# Machine learning predictions improve identification of real-world cancer driver mutations

**DOI:** 10.1101/2024.03.31.587410

**Authors:** Thinh N. Tran, Chris Fong, Karl Pichotta, Anisha Luthra, Ronglai Shen, Yuan Chen, Michele Waters, Susie Kim, Michael F Berger, Gregory Riely, Marc Ladanyi, Debyani Chakravarty, Nikolaus Schultz, Justin Jee

**Affiliations:** Memorial Sloan Kettering Cancer Center

## Abstract

Characterizing and validating which mutations influence development of cancer is challenging. Machine learning has delivered significant advances in protein structure prediction, but its utility for identifying cancer drivers is less explored. We evaluated multiple computational methods for identifying cancer driver alterations. For identifying known drivers, methods incorporating protein structure or functional genomic data outperformed methods trained only on evolutionary data. We further validated VUSs annotated as pathogenic by testing their association with overall survival in two cohorts of patients with non-small cell lung cancer (N=7,965 and 977). “Pathogenic” VUSs in *KEAP1* and *SMARCA4* identified by several methods were associated with worse survival, unlike “benign” VUSs. “Pathogenic” VUSs exhibited mutual exclusivity with known oncogenic alterations at the pathway level, further suggesting biological validity. Despite training primarily on germline, rather than somatic, mutation data, computational predictions contribute to a more comprehensive understanding of tumor genetics as validated by real-world data.

## Main

The majority of somatic tumor mutations are variants of unknown significance (VUSs)^1–3^. In a pan-cancer, multi-institutional cohort of N=160,969 patients with tumor genomic profiling^4^, approximately 80% of somatic mutations detected were VUSs according to an FDA-recognized molecular knowledge database (OncoKB^5^). In some genes with known consequences for survival, such as *KEAP1*^*6*^, 78.8% were VUSs **(Figure 1A)**.

**Figure 1.**
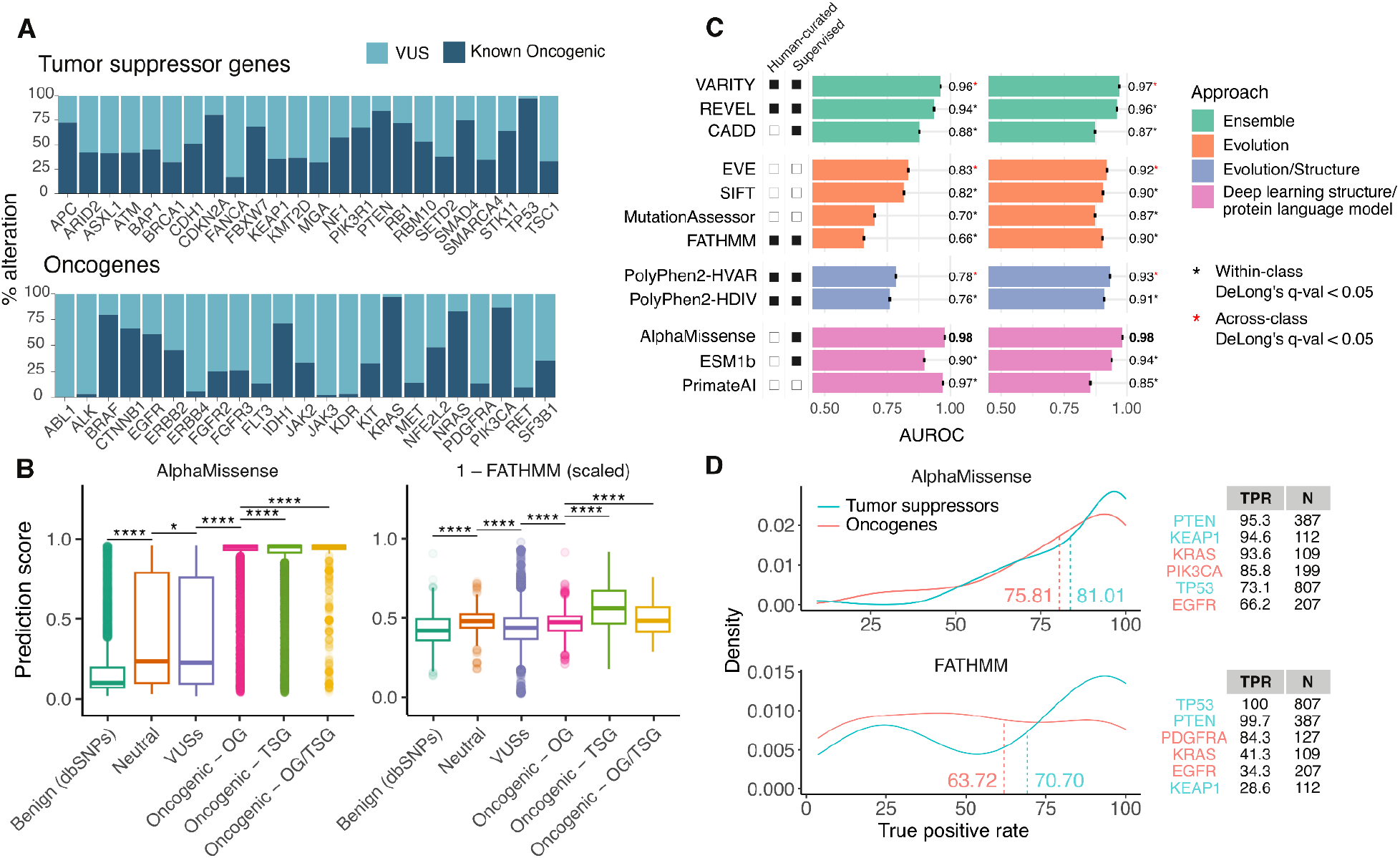
VEPs have variable performance in annotating known oncogenic mutations. A. Frequency of known oncogenic mutations and VUSs in commonly altered oncogenes and tumor suppressor genes in GENIE as annotated by OncoKB. B. Distributions of prediction scores from AlphaMissense and FATHMM from 10,000 non-pathogenic dbSNPs and missense mutations in GENIE v.14-public, broken down by their occurrence in oncogenes (OG), tumor suppressor genes (TSG) or genes that act as both (OG/TSG) at the population level, in which all occurrences of missense mutations are included. For ease of comparison, scores from FATHMM were scaled to the 0-1 range and the difference between 1 and the (scaled) prediction scores was plotted, so that points higher on the y-axis corresponded with higher predicted pathogenicity. Boxplots depict means +-interquartile ranges. C. Bar chart showing AUROC (+/-95%CI) of six variant annotation methods in classifying known oncogenic mutations and non-oncogenic SNPs at the population level. Significant differences between AUROCs were tested using DeLong’s tests and p-values were corrected for multiple hypothesis testing. Within each class of approach, pairwise comparisons were carried out between the best-performing method and all other methods of the same class (significant differences denoted with black asterisks). Pairwise comparisons between the best-performing method of each approach and the best overall method (denoted by bold AUC) were also carried out, with significant differences denoted by red asterisks. D. Density plots showing true positive rates (TPR) of AlphaMissense and FATHMM over all genes. TPR is defined as the number of known oncogenic mutations accurately annotated as pathogenic by AlphaMissense divided by the total number of known oncogenic mutations. TPRs and the number of known oncogenic mutations (N) in select commonly mutated oncogenes and tumor suppressor genes are shown. See Supplemental Appendix for a complete list of TPRs.

Multiple knowledge bases have been developed to annotate pathogenic and actionable mutations^5,7,8^, however, these generally rely on published literature, which is time-consuming to produce and to compile. Computational variant effect predictors (VEPs) may automate variant annotation. In particular, recent tools such as Google DeepMind’s AlphaMissense, which leverage evolutionary, biological and protein structural features combined with high-dimensional machine learning architectures, have gained significant interest^9,10^. However, these VEPs are generally trained to predict germline pathogenic variants, and their utility in identifying somatic mutations that drive diseases such as cancer remains uncharacterized^9,11–15^. The validation of such tools is itself a challenging task; functional assays are labor-intensive and can thus characterize only a limited number of variants^3,16,17^.

Digital health records and widespread tumor genomic profiling offer a means to bypass traditional functional assays and study the impact of tumor mutations directly using patient data^18^. Here, we developed four specific approaches to assess the utility of computational methods for annotating VUSs using real-world patient cancer data: 1) annotating known pathogenic somatic cancer variants as confirmed by OncoKB, which combines literature-confirmed annotations with population-level hotspot identification^19^, 2) identifying VUSs associated with binding regions among proteins with known structures, 3) identifying VUSs associated with overall survival (OS) in patients with lung cancer and 4) identifying VUSs in tumors without other drivers in the same oncogenic pathway. We applied these four approaches to evaluate 11 modern computational methods chosen based on their novelty and demonstrated superior performance in annotating known pathogenic mutations in databases such as ClinVar ^20^ and VariBench ^21^ compared to other methods in the same class^22,23^ (**Table S1**).

We first tested the utility of VEPs in discriminating literature-confirmed or hotspot pathogenic somatic missense variants from benign ones. OncoKB-annotated pathogenic variants in the AACR Project GENIE dataset^4^ served as pathogenic cases while randomly selected missense mutations from the dbSNP Human Variation Sets labeled as having no known medical impact, served as negative controls. Pathogenicity predictions from all 11 methods were generally correlated with each other at both the mutation level, where each unique mutation was counted once, and at the population level, where all occurrences of mutations were counted to reflect actual population frequencies of each mutation **(Figure S2)**.

Benign SNPs had significantly lower scores than oncogenic mutations in all studied methods, demonstrating their ability to distinguish benign versus pathogenic mutations in cancer (**Table S4, Figure 1B, Figure S3**, SA: Predicting Established Pathogenic Variants). Mutations in GENIE annotated as neutral or unknown by OncoKB had significantly higher scores than known benign mutations, suggesting potential unannotated drivers (**Figure 1B**). Across all methods, oncogenic mutations in tumor suppressor genes (TSGs) had higher predicted scores and higher AUROC for correctly annotating oncogenic mutations in TSGs than in oncogenes (OGs), as expected based on previous work^24^.

We found that in general, the ensemble and deep learning-based methods outperformed the evolution-based methods in predicting both TSG and OG oncogenic mutations (**Figure 1C**). AlphaMissense significantly outperformed other deep learning-based methods as well as other best-in-class methods in predicting oncogenic mutations (AUROC of 0.92 and 0.94 for OGs and TSGs at the population level respectively, **Figure 1C**). Among ensemble methods, VARITY and REVEL, both trained on human-curated data, outperformed CADD, which was trained on weak population-derived labels (**Figure 1C**). Among evolution-based methods, EVE, the only unsupervised deep learning method in this class, outperformed others at the population level (AUROC of 0.83 and 0.92 for OGs and TSGs respectively, **Figure 1C**). These results held when mutant alleles were each counted once, irrespective of population frequency **(Figure S4)**. Across all methods, sensitivity was higher in TSGs compared to OGs, but within each gene group, sensitivity further varies at the gene level (**Figure 1D, Table S5**). Overall, these results demonstrate that VEPs were able to identify pathogenic mutations in cancer, with multimodal, deep learning-based methods outperforming methods trained only on mutation frequencies.

We next investigated the ability of VEPs to identify new driver mutations not previously annotated by OncoKB from the large pool of detected VUSs. In particular, we validated the potential functional impact of VUSs labeled as pathogenic by VEPs (“reclassified pathogenic”) in cancer through analyses of their impact on protein binding sites, correlation with patient outcomes, and adherence to expected driver mutual exclusivity patterns.

Pathogenic mutations can alter protein function by disrupting interactions with other proteins and ligands^25^. We probed whether reclassified pathogenic variants were enriched in residues involved in ligand or protein-protein binding (“binding residues”) for proteins with available crystal structures. Mutations affecting binding residues in all genes were significantly more likely to be annotated as oncogenic by OncoKB as expected (Fisher’s test, p-value < 0.001, **Table S6A, Figure 2A**). Mutations occurring at binding residues were universally more likely to be reclassified as pathogenic, whereas non-binding residue mutations were more likely to be reclassified as benign, although the degree of enrichment varied by method (**Figure 2A**). Across all VEPs, VARITY and AlphaMissense achieved the greatest difference in the percent of reclassified VUSs occurring in binding versus non-binding residues (48% and 45%, respectively). This result suggests that the disruption of function at these critical binding residues may contribute to the pathogenic nature of these reclassified pathogenic variants.

**Figure 2.**
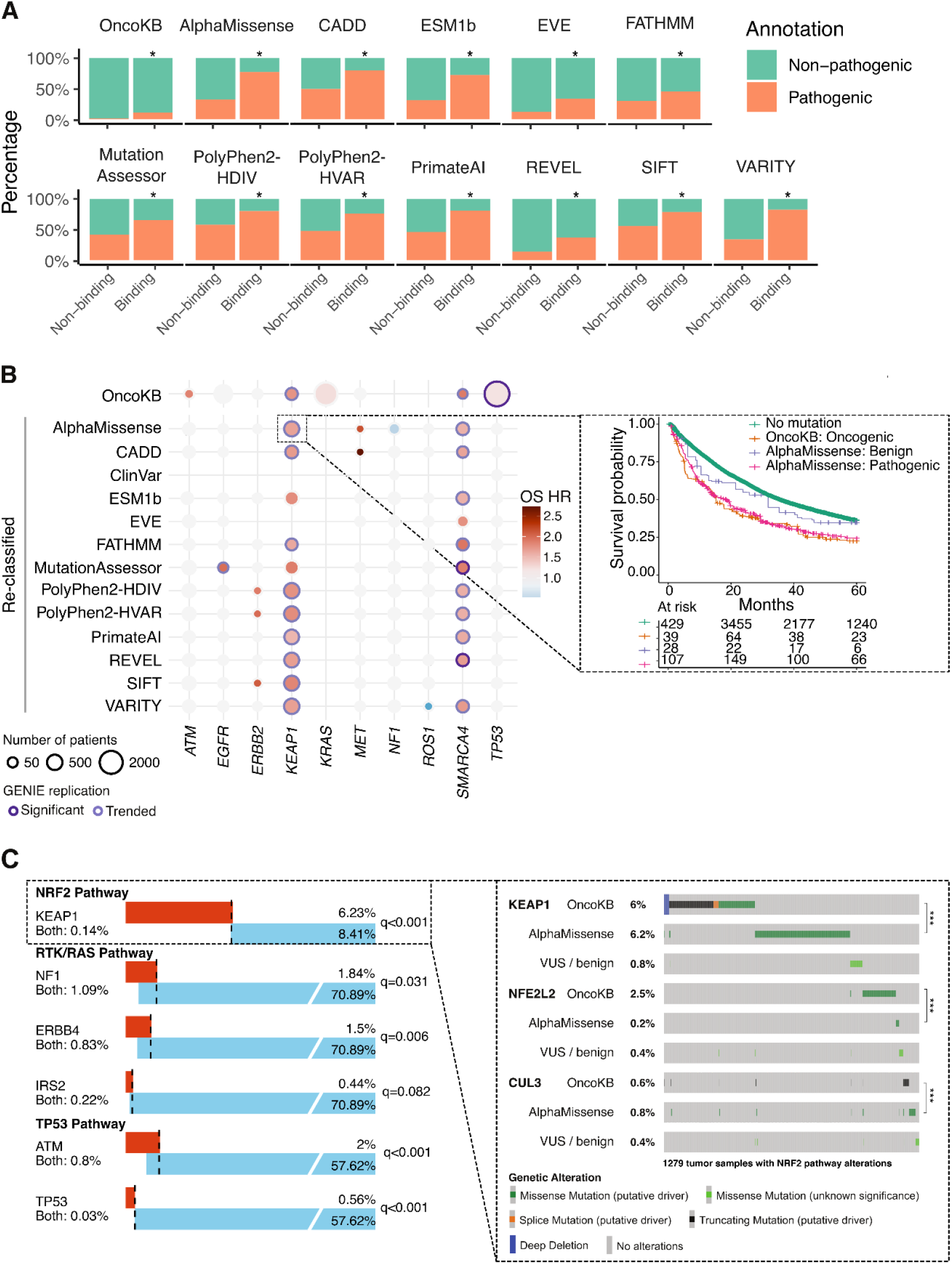
VEPs identify unannotated driver mutations. A. Frequency and annotation of missense mutations occurring at binding residues (either ligand binding or PPI hotspots, see Supplemental Appendix) or non-binding residues of all genes with available binding residue information in GENIE v14-public pan-cancer cohort, Stars denote significance (q-value <= 0.1) in Fisher’s exact tests. OncoKB groups include all missense mutations, whereas variant effect predictor groups only include VUSs. B. Inverse probability treatment weighted overall survival hazard ratios (from time of diagnosis left truncated at time of sequencing) of patients harboring reclassified oncogenic mutations compared to patients without mutation in commonly mutated genes in non-small cell lung cancer (NSCLC). Patients are from MSK-IMPACT NSCLC (N=7,965) and AACR GENIE Biopharma Collaborative NSCLC (N=977) cohorts. Inset: Inverse Probability of Treatment Weighted Kaplan Meier curves comparing overall survival from time of diagnosis left-truncated at time of sequencing of patients based on *KEAP1* mutations annotation in the MSK-IMPACT NSCLC cohort. C. Alteration frequencies and overlap of AlphaMissense reclassified pathogenic mutations (red) with oncogenic alterations of genes in the same pathway (blue) in MSK-IMPACT NSCLC cohort. Inset: Oncoprint of genes in the NRF2 pathway. KEAP1 reclassified mutations, similar to *KEAP1* oncogenic mutations, are mutually exclusive with other oncogenic mutations in *NFE2L2* and *CUL3*. See Supplemental Appendix for details.

To further validate VUS classifications by VEPs using real-world data, we measured the impact of mutations according to classification on OS, focusing on patients with non-small cell lung cancer (NSCLC), the world’s leading cause of cancer mortality^26^ and a cancer type with frequent tumor genomic sequencing. In patients with NSCLC, mutations in multiple genes, including *KRAS, STK11* and *KEAP1*, have been associated with worse OS^27–30^. We investigated whether reclassified pathogenic variants are associated with OS in NSCLC using two cohorts of patients: 7,965 patients with MSK-IMPACT clinical sequencing and 977 non-MSK patients from the GENIE Biopharma Collaborative (BPC) NSCLC cohort^31^. To identify the association between reclassified pathogenic variants and outcome, we stratified patients based on gene-level pathogenicity annotations and compared OS between groups using Cox’s proportional hazard (PH) models, controlling for demographic and clinical covariates by inverse probability of treatment weights (SA: Survival Analysis, **Figure S5**).

Known oncogenic variants^3^ in several genes were associated with worse OS. VUSs in *KEAP1* and *SMARCA4* annotated pathogenic by multiple methods were also associated with worse OS, while those annotated as likely benign were associated with better outcome, suggesting meaningful discrimination among VUSs by computational methods (**Figure 2B, Table S7A**). These findings were consistent in both the MSK-IMPACT and BPC cohorts (**Figure 2B**). The higher OS risks in patients with reclassified pathogenic mutations in these two genes were comparable with the risks associated with known oncogenic mutations, suggesting high specificity for pathogenic variant detection across methods (**Figure S6A, Table S7B**). Conversely, patients with reclassified benign mutations in these genes had some differences in OS risk compared to those with no mutation; for example, CADD, FATHMM, and PrimateAI appeared to have imperfect sensitivity for pathogenic *KEAP1* mutations (**Figure S6B, Table S7B**).

Concurrent mutations in certain gene combinations may worsen survival in an additive manner, as is known for *STK11* and *KEAP1* in NSCLC^28^ and observed in the MSK-IMPACT NSCLC cohort (**Figure S9B**). To test whether reclassified pathogenic variants in *STK11* and *KEAP1* were similarly associated with worse survival, we compared OS of patients with double *KEAP1* and *STK11* reclassified mutations with patients with single reclassified pathogenic mutation and patients without any mutation in these two genes. We found that patients with tumors harboring both *KEAP1* and *STK11* AlphaMissense mutations had worse OS compared to those with reclassified pathogenic mutations in either genes, as well as those without mutation (**Figure S9A**). This result further suggests that the *KEAP1* and *STK11* reclassified variants from AlphaMissense follow expected patterns of additive prognostic significance, suggesting biologic validity. In summary, VEP annotations suggested several VUSs with biological activity that were confirmed by association with OS in NSCLC.

Oncogenic mutations in genes within the same oncogenic signaling pathway tend to not co-occur in the same patient due to functional redundancy^32,33^. In NSCLC, oncogenic mutations in the RTK/RAS, NRF1 and TP53 pathways have been shown to exhibit mutual exclusivity^33^. To demonstrate that reclassified pathogenic mutations have comparable cancer-driving effect on a pathway as known oncogenic mutations, we aimed to identify whether reclassified pathogenic mutations were mutually exclusive with other known oncogenic mutations in these three pathways within the MSK-IMPACT NSCLC data using two-sided Fisher’s exact tests. All methods were able to identify VUSs that exhibit mutual exclusivity with other oncogenic mutations in all three pathways (**Figure S10A**).

Within the NRF2 pathway, *KEAP1* VUSs reclassified as pathogenic by any method were mutually exclusive with oncogenic mutations in *KEAP1, NFE2L2* and *CUL3* (**Figure 2C**), whereas *KEAP1* VUSs reclassified as benign by VEPs except for CADD, ESM1b, FATHMM and MutationAssessor tended to co-occur with other oncogenic mutations in the pathway (**Figure S10B**). Similarly, all VEPs except for MutationAssessor were able to identify reclassified pathogenic mutations in *ATM* and *TP53* that are mutually exclusive with other oncogenic mutations in the TP53 pathway, whereas reclassified benign mutations in these genes were not mutually exclusive with other drivers in the pathway (**Figure 2C, Figure S10**).

In the RTK/RAS pathway, reclassified pathogenic mutations in *NF1, ERBB4*, and *IRS2*^*34*^ identified by AlphaMissense were mutually exclusive with other oncogenic mutations (**Figure 2C**). Reclassified benign mutations in *ERBB4* and *IRS2* were also mutually exclusive with known oncogenic mutations (**Figure S10B**), indicating potential drivers requiring additional annotation. *ERBB4* and *IRS2* pathogenic mutations classified by AlphaMissense were frequent in high TMB samples, while RTK/RAS oncogenic mutations were more common in TMB-low samples (**Figure S11A**). Logistic regressions, controlling for TMB status, revealed independent mutual exclusivity between RTK/RAS oncogenic mutations and reclassified pathogenic mutations in *ERBB4* (**Figure S11B**). A negative association was observed between *IRS2* pathogenic mutations and RTK/RAS oncogenic mutations, although not statistically significant due to limited sample size (**Figure S11C**). Similar patterns were seen with reclassified benign mutations, suggesting potential unannotated drivers in *ERBB4* and *IRS2* (**Figure S11C**). More details about the mutational pattern of RTK/RAS pathway and TMB status of samples with *ERBB4* and *IRS2* mutations are summarized in **Table S9**. In summary, analysis of mutational patterns showed that reclassified pathogenic mutations followed expected patterns within oncogenic pathways, offering a potential benchmark for VEP performance, although greater sample size may be required to fully explore less common driver classes.

Our study has limitations. Our VUS quantification across institutions comes from multiple contributing cancers with their own sequencing pipelines, the majority of which are tumor-only sequencing. Even though all data went through germline SNP filtering pipeline before public release, it is possible that there remained private SNPs in the data, which may artificially inflate the number of more easily characterizable VUSs in a given dataset, although this would reflect a clinical reality of such SNPs appearing in tumor-only sequencing assays. The MSK-IMPACT and GENIE BPC cohorts, though richly annotated, may not be sufficiently powered at the present to discover rare driver variants in less commonly mutated genes or assert the association between these putative drivers and outcomes.

Overall, the findings underscore the potential of VEPs in identifying driver mutations in cancer and find them successful at several benchmarks based on real-world data for predicting cancer driver variants. We expect that VEPs will continue to improve over time, particularly with regard to cancer driver prediction; that real-world datasets fueling these analyses will continue to grow; and that a growing number of molecularly targeted therapies will allow for examination of not prognostic but also predictive value for identified genomic targets, together suggesting synergistic means by which data and computation can improve the lives of patients with cancer.

