## Supplemental Appendix for "Machine learning predictions improve identification of real-world cancer driver mutations"

### Supplementary Appendix for “Machine learning predictions improve identification of real-world cancer driver mutations”

#### Methods

##### Patients and Data Collection

This study analyzed patients with tumor genomic sequencing from two sources: The MSK-IMPACT cohort and the AACR Project GENIE cohort, which includes patients from the MSK-IMPACT cohort.

MSK-IMPACT

The MSK-IMPACT cohort comprised patients at Memorial Sloan Kettering (New York, NY), an academic cancer hospital with tumor genomic sequencing using MSK-IMPACT, an FDA-authorized tumor genomic profiling assay, which uses matched white blood cell sequencing to filter clonal hematopoietic and germline variants. All MSK patients were enrolled as part of a prospective sequencing protocol (NCT01775072). The study was independently approved by the Institutional Review Board of each site. Patients provided written, informed consent and were enrolled in a continuous, nonrandom fashion. Data here is from a February 15, 2023 snapshot, consisting of 11,649 samples from 7,965 patients with non-small cell lung cancer (NSCLC).

For patients in the MSK-IMPACT cohort, demographic and clinical information, including tumor stage, age, sex, race and histology were retrieved from the electronic health records database. Smoking history, prior treatment and metastatic events were abstracted from clinical notes using previously validated natural language processing methods [1–4]. Tumor mutation burden per sample was calculated as the total number of nonsynonymous mutations divided by the actual number of bases analyzed, and samples with TMB >= 10 mut/Mb were defined as TMB-high. MSI-H status was defined for each sample by an MSIsensor score >10 [5].

GENIE

Details of the AACR Project GENIE cohort have been published previously[6]. In short, the pan-cancer registry contains genomic and clinical data from 11 international institutions. In this study, we analyzed data from the v.14-public release, which consists of genomic data for 183,302 tumors from 160,965 patients. For genomic landscape analyses involving gene-level counts, only patients with tumor sequencing panels including a given gene of interest were included in the respective analysis. The number of patients with each gene sequenced is given in **Table S10**.

GENIE BPC

Genomic and clinical data for a subset of patients with non-small cell lung cancer in the AACR GENIE cohort have been recently published as part of the AACR GENIE Biopharma Collaborative (BPC)[7]. Out of 1,846 patients from four contributing institutions in the cohort, we included all 977 patients from Dana-Farber Cancer Institute, Vanderbilt-Ingram Cancer Center and Princess Margaret Cancer Centre-University Health Network with single-primary NSCLC in our analysis cohort. All MSK patients were included in the MSK-IMPACT cohort. For survival analyses involving gene-level cohorts, only patients with tumor sequencing panels including a given gene of interest were included in the respective analysis.

**Genomic Landscape**

Genomic data, including mutational calls, copy number alteration and structural variant data, for the GENIE v.14-public cohort were obtained from Synapse. All genomic alterations were annotated with OncoKB version 4.2 (release date February 10, 2023). Genes were labeled as oncogenes or tumor suppressor genes using the OncoKB Cancer Gene List (updated October 2, 2023). For genomic landscape analyses involving gene-level counts, only patients with tumor sequencing panels including a given gene of interest were included in the respective analysis.

**Non-oncogenic Variants**

Non-oncogenic missense mutations were randomly selected from the dbSNP Human Variation Sets build 150 (April 2017) labeled as having “no known medical impact.” This dataset includes variants with germline minor allele frequency of ≥ 0.01 and no records of clinical phenotypes in ClinVar. We randomly selected 10,000 variants and annotated them with the same eight methods as described below. These non-oncogenic variants served as the negative class in subsequent receiver operating characteristic curve analyses.

**Predicting Established Pathogenic Variants**

We evaluated the performance of six variant function prediction methods and one human variant archive for recapitulating known oncogenic cancer variants as annotated by OncoKB, the first FDA-recognized somatic molecular knowledge database for this purpose. Methods were chosen to be included based on their novelty (including methods with recent release dates e.g. AlphaMissense, PrimateAI and ESM1b, as well as novel methodology to an old approach e.g. EVE’s deep generative model to predict pathogenicity based on evolutionary conservation), superior performance compared to other methods in the same class (e.g. REVEL and CADD outperformed other VEPs in multiple comparison studies[8,9], while VARITY_R_LOO’s performance was better than 12 other VEPs and comparable to AlphaMissense in certain evaluation tasks[10]), specific relevance to cancer (FATHMM implements cancer-specific pathogenicity weight[11] and MutationAssessor previously demonstrated utility in annotating cancer variants[12]), as well as historical significance and popularity (SIFT and PolyPhen2 are among the earliest VEPs and have the highest number of citations to date[13]). A brief description of the methods/database is presented in **Table S1**.

**Annotation schema**

OncoKB annotations were performed with version 4.2 (released February 10, 2023). Prediction scores for all VEPs and ClinVar were obtained from dbNSFP[14] v4.6 (released February 18, 2024). Categorization of variants into pathogenic, non-pathogenic or uncertain, obtained by imposing predetermined cutoff thresholds on the predicted functional scores, was also provided by AlphaMissense, FATHMM, MutationAssessor, PolyPhen2 and SIFT. For VEPs in this category, we used off-the-self classifications provided by these methods. The rest of the evaluated VEPs required additional steps to determine the appropriate variant classification from predicted scores.

First, EVE provides variant classifications at different degrees of uncertainty, aiming to maximize accuracy by excluding variants that the model is uncertain about. For example, a Class25 EVE classification means that 25% of the most uncertain variants were excluded when making predictions[15]. To identify the degree of uncertainty that would maximize accuracy in our data, we compared the AUROCs for classifying known oncogenic mutations from benign dbSNPs and selected the uncertainty threshold that resulted in the highest AUROC. Finally, for VEPs that provided prediction scores but not off-the-shelf classification or recommended score cutoffs for classifications, including CADD, ESM1b, PrimateAI, REVEL and VARITY, we manually identified cutoff thresholds to classify pathogenic and non-pathogenic mutations. To identify cutoff, for each method, we set up a binary classification task in which known oncogenic mutations from the dataset represented the positive class, and the benign dbSNPs represented the negative class. We then calculated the sum of sensitivity and specificity of each method at different thresholds and identified the optimal cutpoint as the threshold where this sum was maximized. In cases where more than one cutpoint were found, we used their median as the final threshold.

After cutoffs were determined, we applied them to the VEPs' predicted scores to stratify variants into categories. For CADD, PrimateAI, REVEL and VARITY_R_LOO, variants with scores higher than or equal to the determined thresholds were classified as pathogenic and vice versa. For ESM1b, variants with scores smaller than or equal to the determined thresholds were classified as pathogenic.

We repeated this process for all datasets, resulting in data-specific cutoff and EVE uncertainty thresholds. The cutoffs used are summarized in **Table S2**.

**Receiver operating curves (ROCs)**

We performed two ROC analyses, 1. Weighting all positive variants equally and 2. To better understand method performance in a manner that reflects actual population levels of a given mutation, sampling positive variants from the GENIE cohort, i.e. appearing proportionally to their frequency in a real-world population. In the first analysis, mutation-level ROC for each method was constructed using all 8,033 missense mutations as the positive class and 10,000 dbSNPs as the negative class. In the second analysis, population-level ROCs were constructed using all occurrences of missense mutations in the GENIE v14-public cohort as the positive class (N=246,401) and 10,000 dbSNPs as the negative class. The non-oncogenic mutations were upsampled to match the number of oncogenic mutations in the positive class, resulting in a balanced N=246,401 for each class. Area under the curve and 95% confidence interval were calculated for each curve.

**Ligand binding and protein-protein interaction residues analysis**

Residues involved in binding ligands, including small molecules, peptides, DNA and RNA, were retrieved from BioLiP2[16], a curated database of biologically relevant protein-ligand interactions. Residues important in protein-protein interactions (PPI), termed PPI hotspots and defined as residues whose replacements decrease the binding free energy significantly, were retrieved from PPI-HotspotDB[17]. We grouped missense mutations in either the GENIE v14-public or MSK-IMPACT NSCLC cohort into binding residues (including ligand-binding residues and PPI hotspots) versus non-binding residues. Fisher's exact tests were performed to test the enrichment of mutations occurring at binding residues for being reclassified as pathogenic by different methods.

##### Pathway Analysis

*Mutual exclusivity test*

Canonical oncogenic pathway level analyses were conducted using curated pathway templates as previously reported[18].

We aimed to identify whether reclassified pathogenic mutations are mutually exclusive with other known oncogenic mutations, which demonstrates that reclassified pathogenic mutations have comparably pathogenic effect on a pathway as known oncogenic mutations. To this end, we calculated one versus all mutual exclusivity for each gene for patients with NSCLC in the MSK-IMPACT cohort. For each test, we first set up a 2x2 contingency table with two variables: the number of patients carrying reclassified pathogenic mutations in that gene, and the number of patients carrying oncogenic mutations in all genes within the same pathway. A two-sided Fisher's exact test was applied to the contingency table to test for mutual exclusivity.

An example contingency table used to test for pathway mutual exclusivity between KEAP1 and all genes in the NRF2 pathway, including KEAP1, CUL3 and NFE2L2, is below:

| | | **Mutations in KEAP1** | |
| --- | --- | --- | --- |
| | | reclassified pathogenic | VUSs/no mutation |
| **Mutations in KEAP1, CUL3 and NFE2L2** | Known oncogenic | | |
| | VUSs/no mutation | | |

In particular, this table is used in a Fisher's exact test for mutual exclusivity between KEAP1 and all genes in the NRF2 pathway. Two other tests were set up to test for mutual exclusivity of NFE2L2 and CUL3 with oncogenic mutations in NRF2 pathway genes.

The procedure was repeated for all genes present in a given pathway, and for all eight methods. P-values were adjusted for multiple hypothesis testing using the Benjamini-Hochberg procedure. Tests with a logOR $< 0$ and adjusted p-value $<= 0.1$ were considered significant for mutual exclusivity. The pathway-level one versus all mutual exclusivity rate was then calculated for each pathway by dividing the total number of significant mutually exclusive tests by the number of genes in the pathway.

As negative controls, we tested for mutual exclusivity between reclassified benign mutations and known oncogenic mutations in all genes in a given pathway using the same procedure.

*The role of tumor mutational burden in observed mutual exclusivity*

To identify whether ERBB4 and IRS2 AlphaMissense reclassified pathogenic mutations are mutually exclusive with RTK/RAS oncogenic mutations independent of TMB-high status, we performed a logistic regression:

RTK/RAS oncogenic mutations ~ AlphaMissense mutations + TMB-H status

Where

*RTK/RAS oncogenic mutations* is 1 if a sample has any oncogenic mutations in RTK/RAS pathway genes, 0 otherwise

*AlphaMissense mutations* is 1 if a sample has an AlphaMissense reclassified pathogenic mutation in a gene of interest (ERBB4 or IRS2), 0 otherwise

*TMB-H* is 1 if a sample has TMB >= 10 mut/Mb, 0 otherwise

Two independent regressions were run for *ERBB4* and *IRS2*.

**Survival Analysis**

To test the association of gene-level pathogenicity annotations with overall survival we performed a series of Cox proportional hazards (PH) models from time of diagnosis to time of death or last follow-up, left truncated at time of cohort entry (tissue sequencing). For patients with multiple sequencing events, the first was used as the time of cohort entry. To adjust for confounding variables between comparison groups, inverse probability of treatment weights (IPTW) were calculated using covariate values at baseline, including tumor stage, age, sex, race, histology, smoking history, tumor mutational burden (TMB), microsatellite instability status (MSI), prior treatment and metastatic sites if any, before fitting Cox PH models. An example plot of standardized mean differences in covariates before and after IPTW matching to demonstrate how IPTW helps achieve balance in covariates between comparison groups is presented in Figure S9.

Hazard ratios, 95% confidence interval and p-values for gene-level associations between a "pathogenic" alteration vs no alteration were computed. In the MSK-IMPACT cohort, all genes altered in >2% of the cohort were considered. For each gene of interest, only patients with tumor sequencing panels including a given gene in the target region were included in the analysis. The following gene mutation annotation schema was used: For OncoKB, any alterations annotated as oncogenic or likely oncogenic were used. For all other databases, any pathogenic alteration considered a variant of unknown significance in OncoKB (i.e. a "reclassified" pathogenic alteration) was used. q-values were computed using the Benjamini-Hochberg method; a false discovery rate 0.1 across all comparisons described in this analysis was used to determine statistical significance according to a prespecified statistical analysis plan (see 12-245 Appendix C Project Plan). The BPC cohort was used as a confirmation dataset in which only significant associations from the MSK-IMPACT analysis were tested; q-values were computed similarly but for only the number of hypotheses tested in the BPC.

*Kaplan-Meier curves*
Weighted Kaplan-Meier (KM) curves were constructed to further examine the relationship between gene-level pathogenicity annotations and overall survival for genes with significant hazard ratios in univariate weighted Cox PH models. Similar to the Cox PH regressions, KM curves were calculated from time of diagnosis to time of death or last follow-up, left truncated at time of cohort entry (tissue sequencing). Patients were stratified based on the presence of OncoKB oncogenic, ‘reclassified’ pathogenic, ‘reclassified’ benign mutations or without any mutation in each gene of interest. Only strata with ≥ 10 patients were included. KM curves were weighted using the same IPTWs used for the corresponding Cox PH regression.

*STK11/KEAP1 concurrent mutation analysis*
To test whether AlphaMissense reclassified pathogenic variants in *STK11* and *KEAP1* were similarly associated with worse survival, we compared overall survival of patients with double *KEAP1* and *STK11* reclassified mutations with patients with single reclassified mutation and patients without any mutation in these two genes. Patients with reclassified pathogenic mutations in both *KEAP1* and *STK11* are labeled as KEAP1/STK11 double mutant, while patients carrying only reclassified pathogenic mutations in either genes are labeled as single mutant for the respective genes. Patients carrying oncogenic mutations in either genes are excluded from this analysis. Weighted KM curves were constructed as described above to compare overall survival of patients in these four strata.

All statistical analyses were performed in R version 4.2.2 (2022-10-21).

#### Supplementary Figures

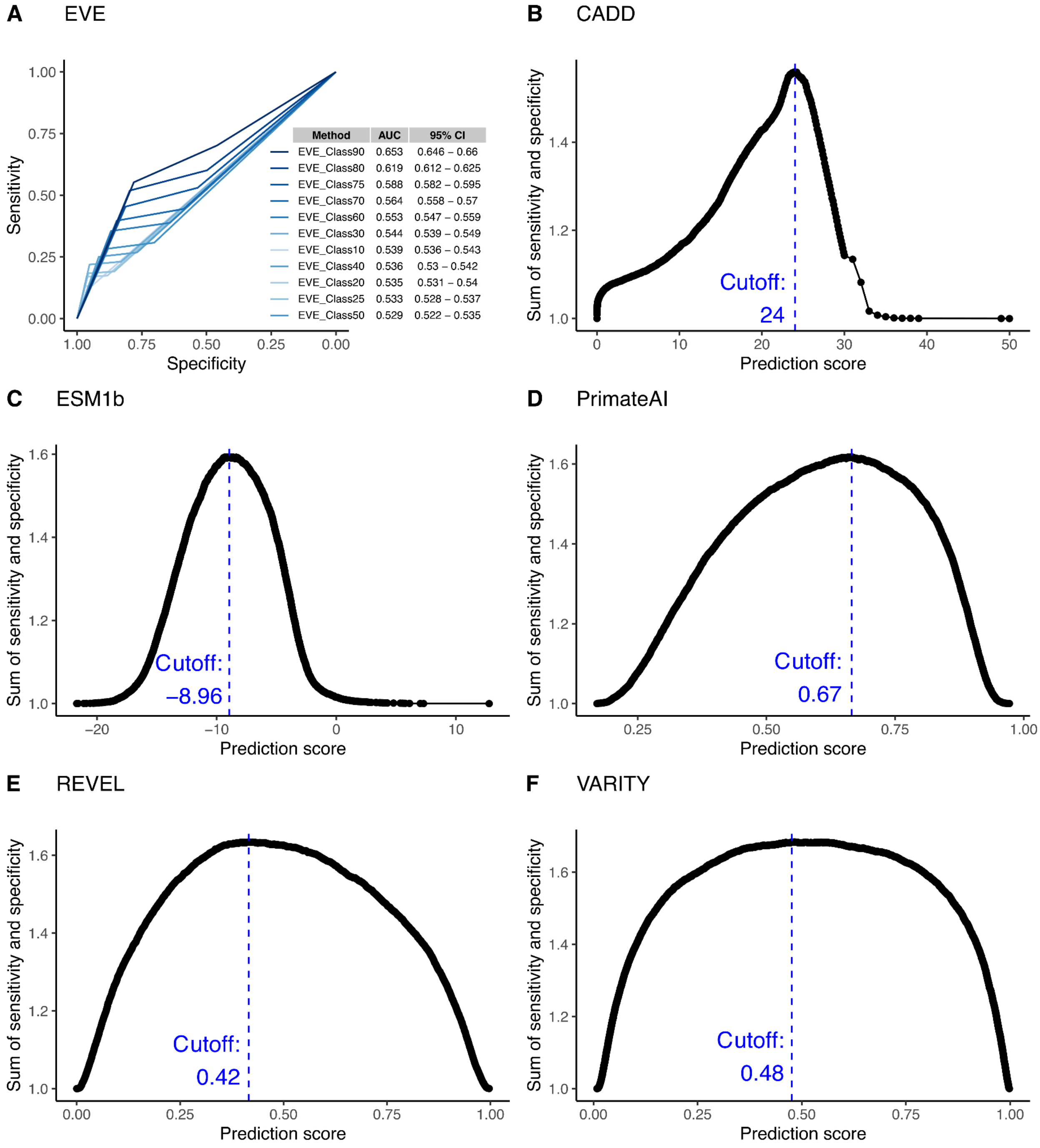


**Figure S1. Determination of CADD score cutoff for pathogenic mutations**

The CADD score cutoff threshold for pathogenic mutations was determined by optimizing the sum of sensitivity and specificity in classifying oncogenic mutations and non-oncogenic dbSNPs.

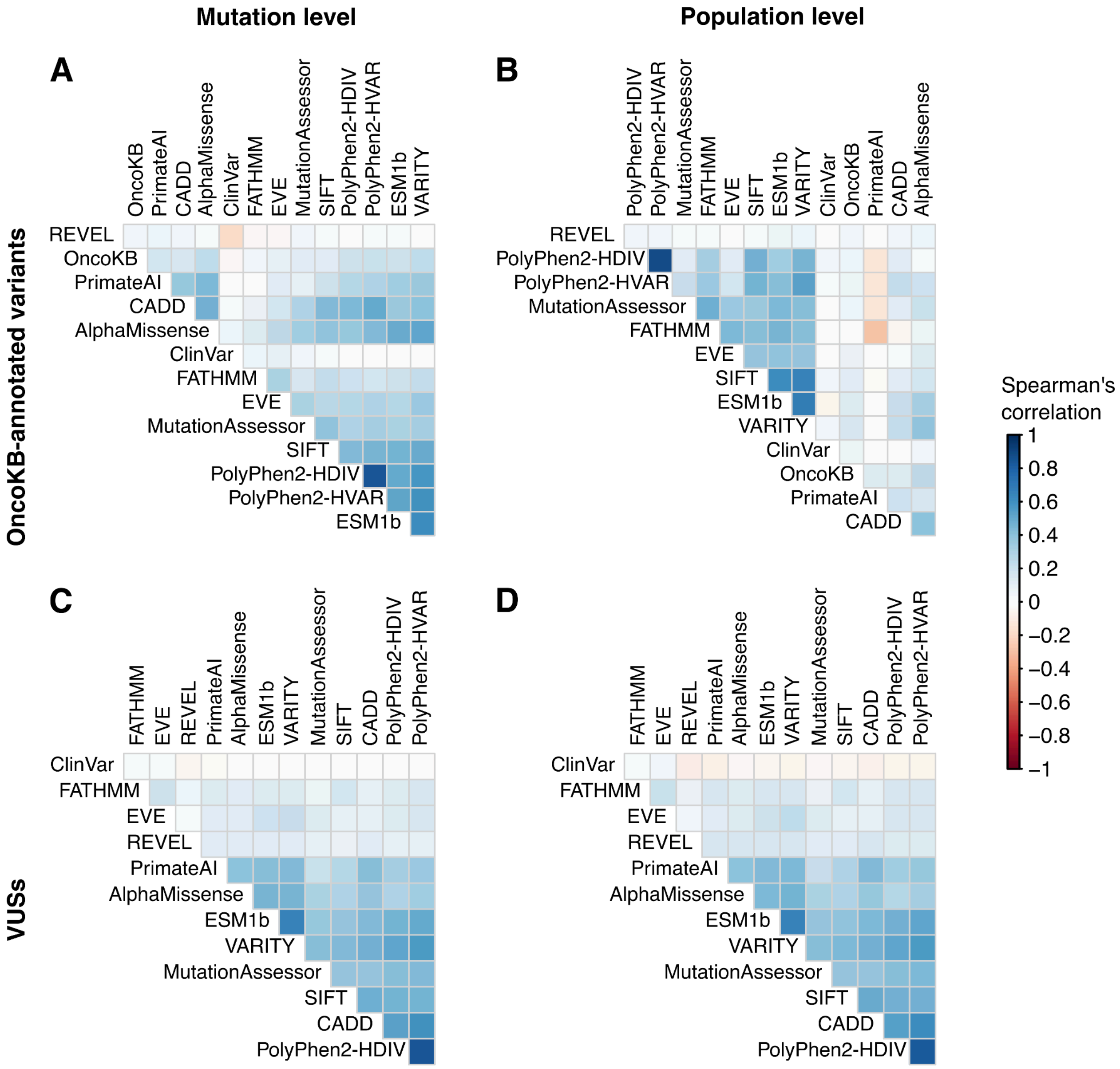


**Figure S2. Concordance heatmap of mutation annotations GENIE**

Spearman's correlation was used to calculate the correlation between the predicted classification by each method (non-pathogenic or pathogenic, as per Supplemental Methods) and as annotated by OncoKB (unknown or oncogenic). Tile colors depict Spearman's correlation coefficients.

A. Correlation at the mutation level, in which each unique mutation is included once.
B. Correlation at the population level, in which all occurrences of missense mutations are included.

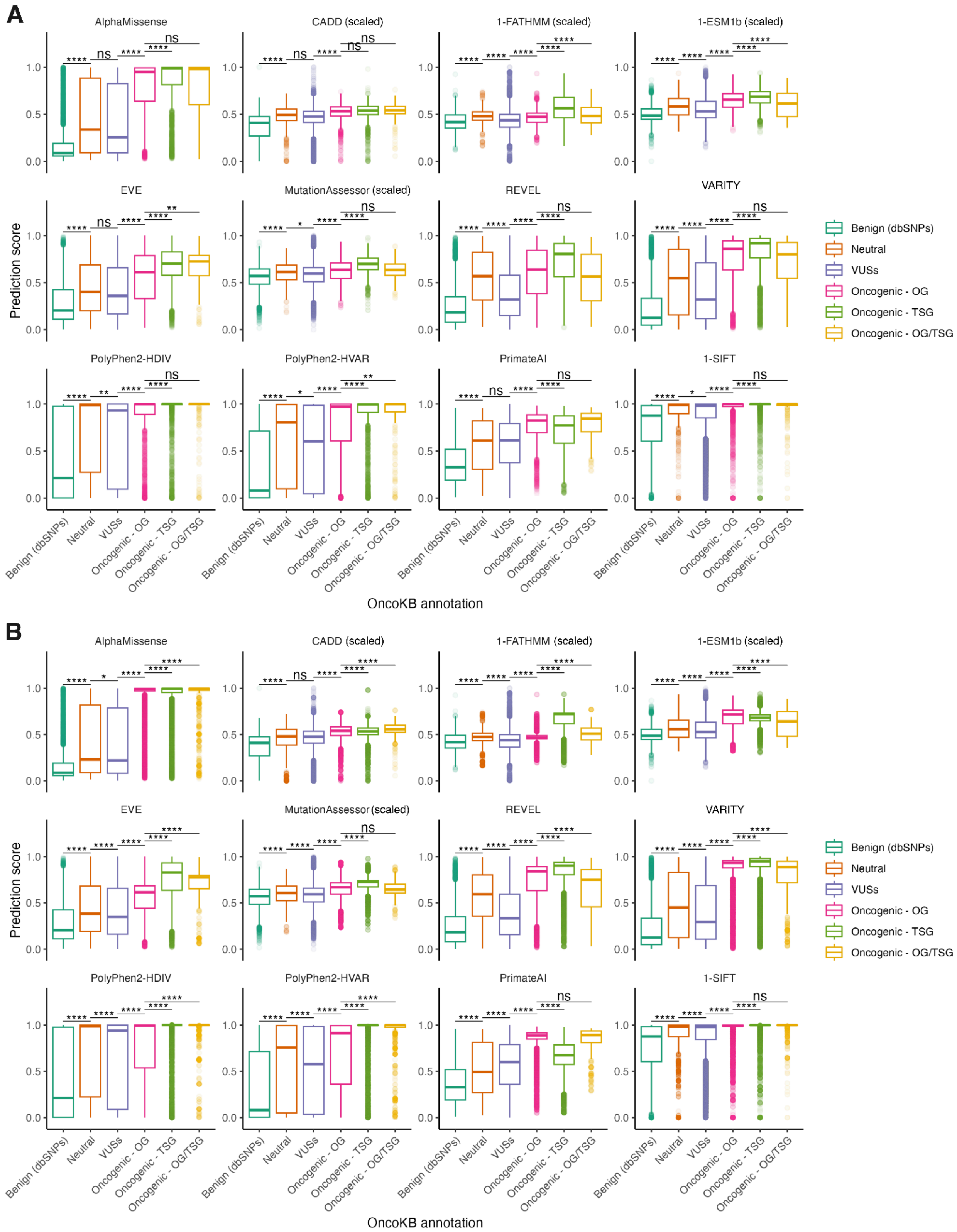


**Figure S3. Prediction scores of missense mutations and benign dbSNPs**
Distribution of prediction scores from benign dbSNPs and missense mutations in GENIE v.14-public, broken down by their occurrence in oncogenes (OG), tumor suppressor genes

(TSG) or genes that act as both (OG/TSG). For most VEPs, scores are in the range of 0-1, with higher scores suggest higher predicted pathogenicity. For ease of comparison, scores from methods with different ranges, including CADD, MutationAssessor, FATHMM and ESM1b, were scaled to the 0-1 range. For methods where lower scores indicated higher pathogenicity, the difference between 1 and the (scaled) prediction scores was plotted. Boxplots depict means +- interquartile ranges.

Pairwise differences in means between groups were tested for significance using Tukey's range test and p-values were corrected for multiple hypothesis testing using the Benjamini-Hochberg method. Asterisks on brackets depict significance levels of q-value.

A. Score distributions at the mutation level, in which each unique mutation is included once.
B. Score distributions at the population level, in which all occurrences of missense mutations are included.

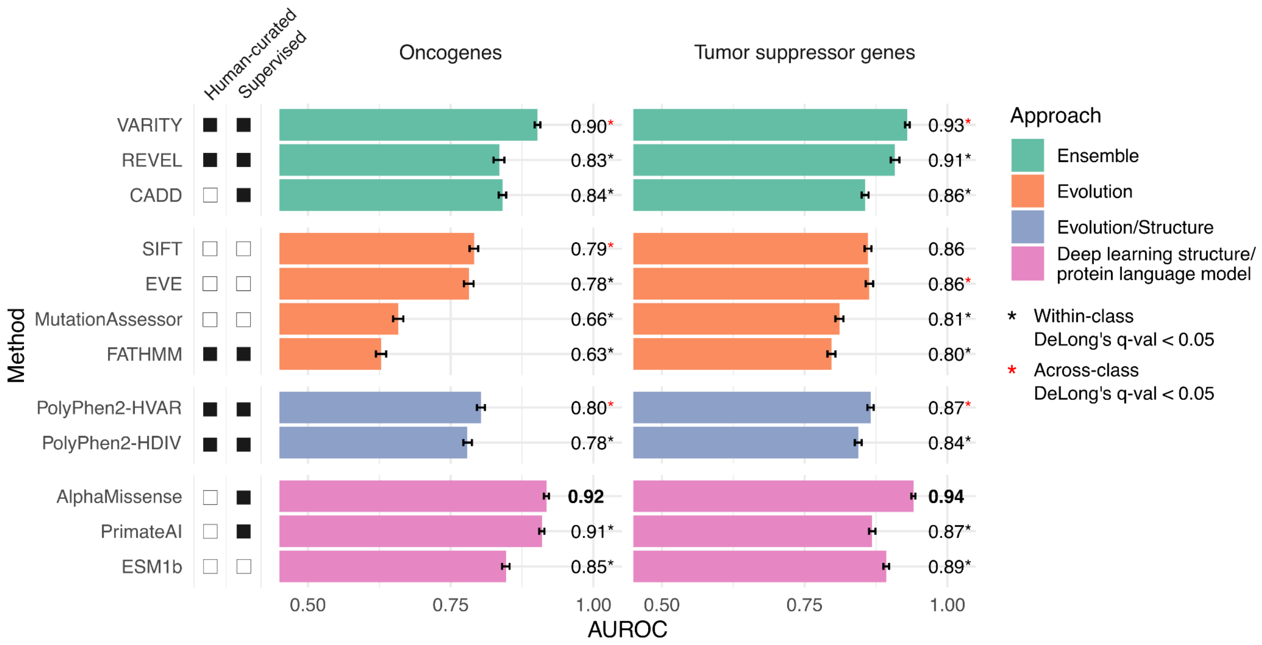


**Figure S4. Mutation-level evaluation of VEP performance in identifying pathogenic cancer mutations.**

A. Distributions of prediction scores from 10,000 non-pathogenic dbSNPs and missense mutations in GENIE v.14-public, broken down by their occurrence in oncogenes (OG), tumor suppressor genes (TSG) or genes that act as both (OG/TSG) at the mutation level, in which each unique missense mutation is counted once. Each gray point on the plot depicts an occurrence of a mutation. Boxplots depict means +- interquartile ranges.

B. Receiver operating curves showing performance of six variant annotation methods in classifying known oncogenic mutations and non-oncogenic SNPs at the mutation level.

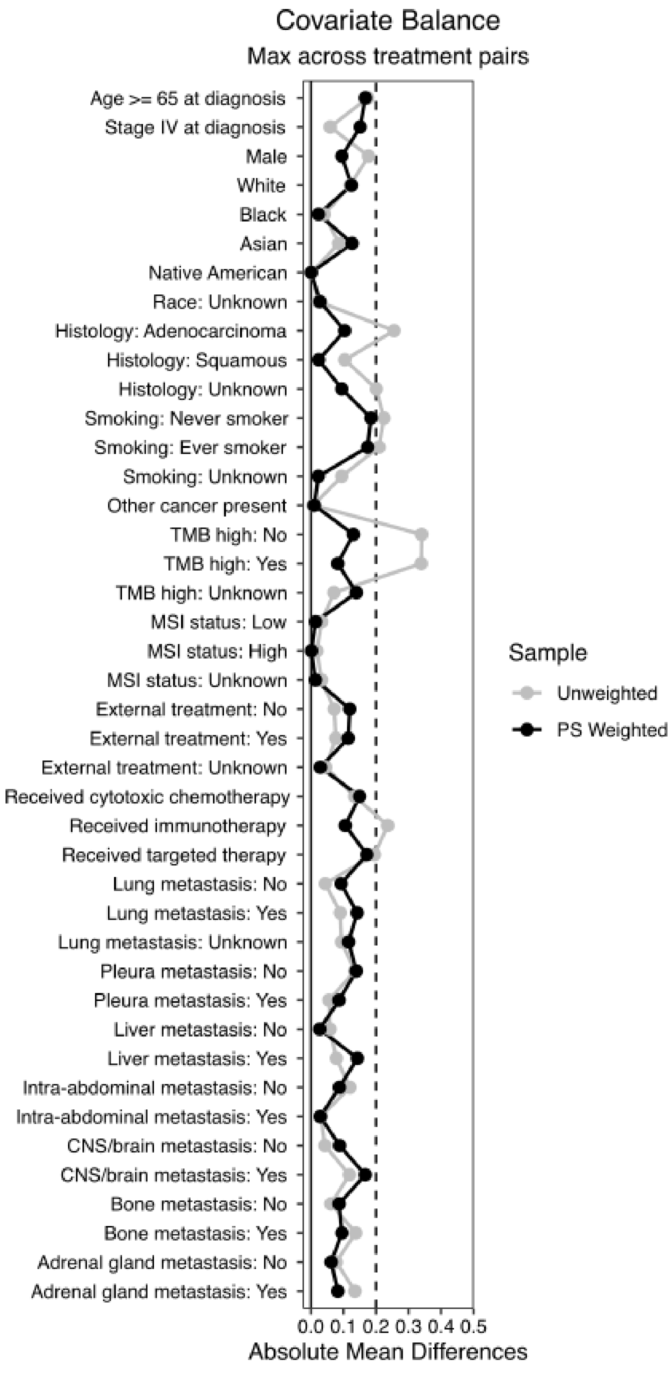


**Figure S5. Balance of covariates between patient strata using inverse probability of treatment scores**

Absolute mean differences between clinical and demographic covariates in four groups of patients stratified by KEAP1 mutational status, including  No KEAP1 mutation, KEAP1 - Oncogenic mutations, KEAP1 - reclassified pathogenic mutations by AlphaMissense, KEAP1 - Reclassified benign mutations by AlphaMissense before and after inverse probability of treatment score weighting (IPTW). IPTW helps achieve balance in variables with large imbalance between groups, such as TMB high status.

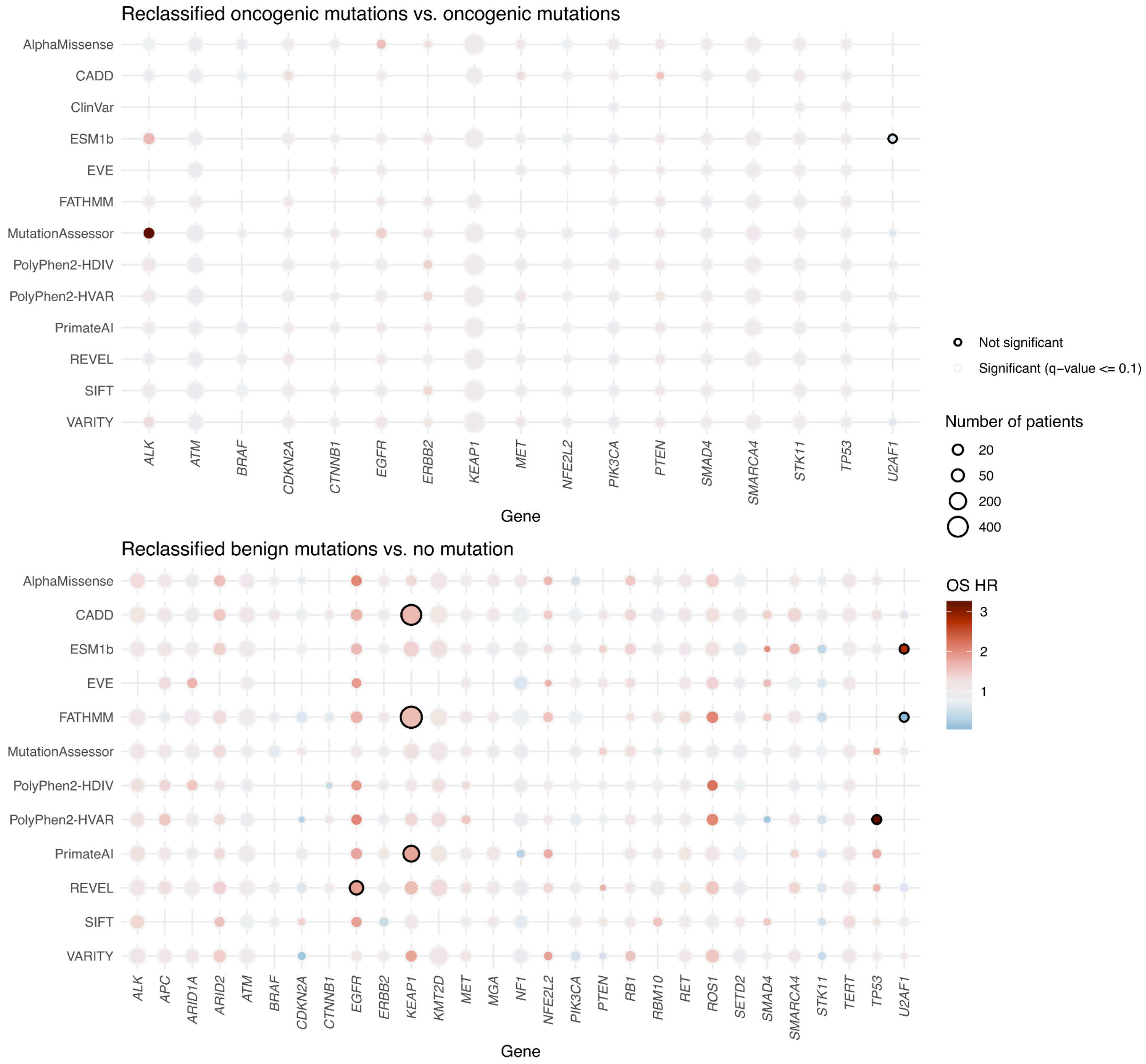


**Figure S6. Cox proportional hazards coefficients of reclassified mutations versus known oncogenic mutations or no mutation**

A. Overall survival hazard ratios of patients with reclassified pathogenic mutations compared to patients with oncogenic mutations were obtained from weighted Cox proportional hazard models.

B. Overall survival hazard ratios of patients with reclassified benign mutations compared to patients with no mutation in a given gene were obtained from weighted Cox proportional hazard models.

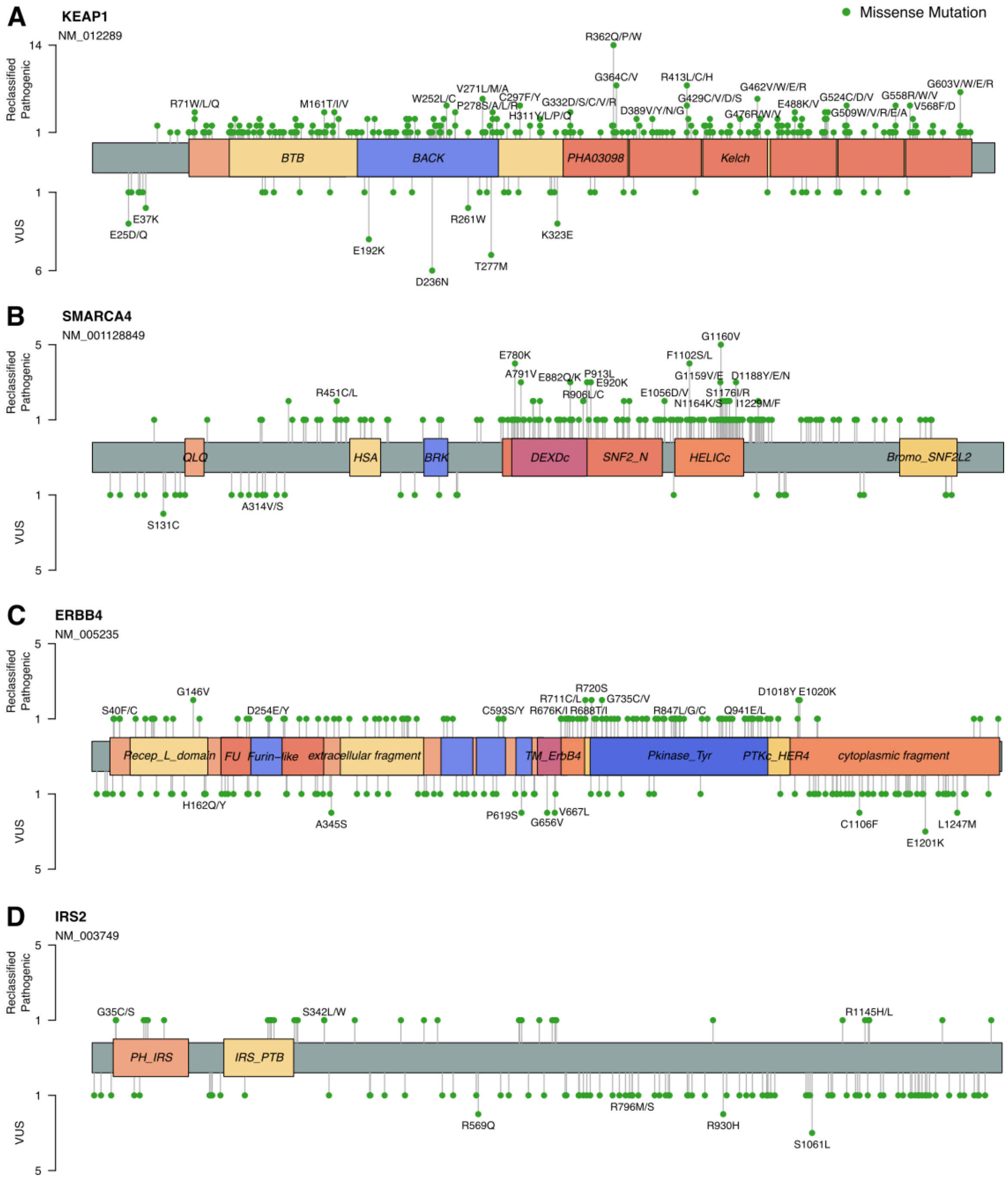


**Figure S7. Lollipop plots of *KEAP1* VUSs reclassified as pathogenic or benign by AlphaMissense**

Amino acid-level changes resulting from AlphaMissense-reclassified pathogenic mutations in in the MSK-IMPACT NSCLC cohort are plotted for:

A. *KEAP1*
B. *SMARCA4*
C. *ERBB4*
D. *IRS2*

The most commonly mutated residues are labeled.

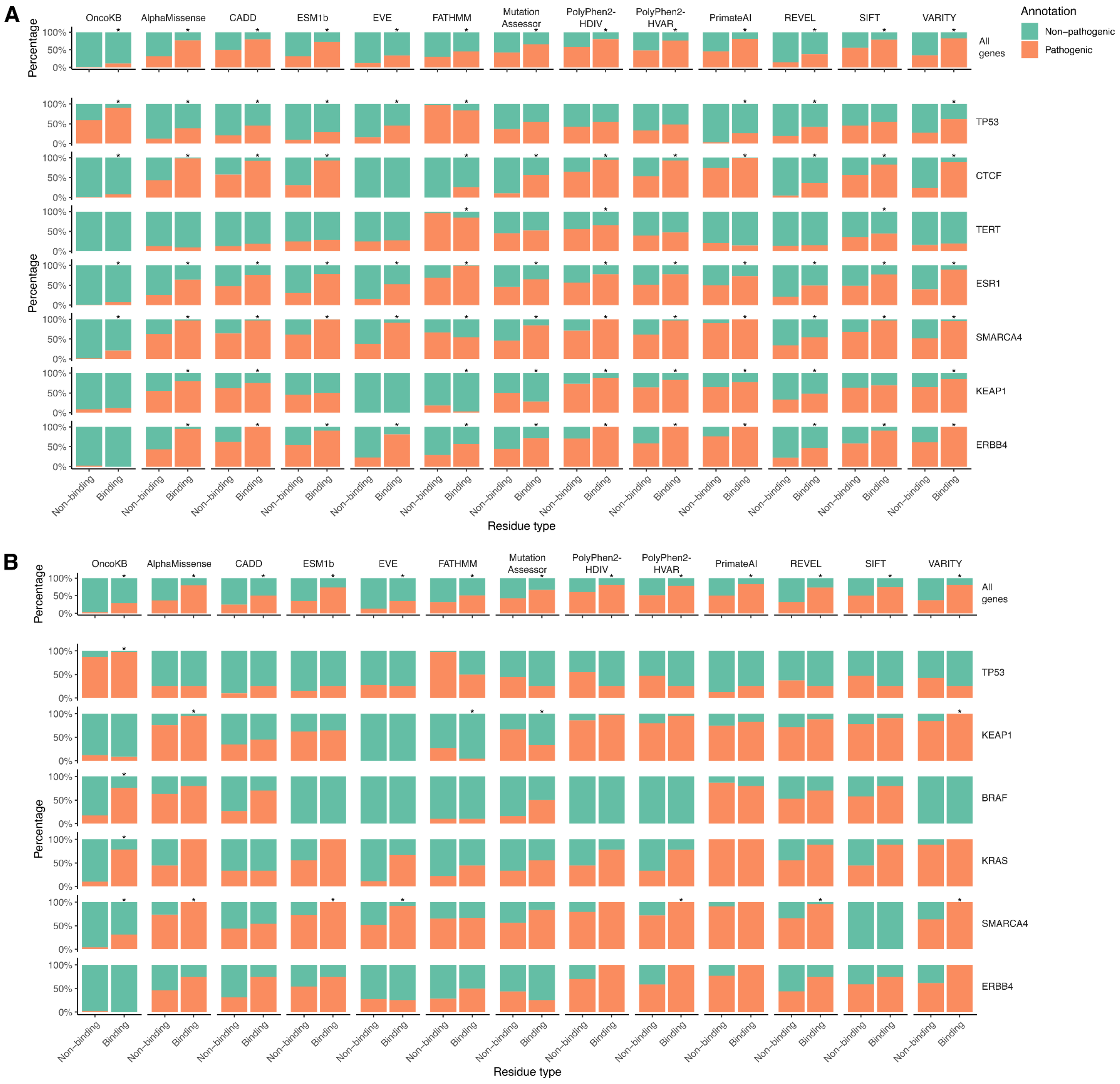


**Figure S8. Reclassified pathogenic mutations occur at ligand-binding residues and protein-protein interaction (PPI) hotspots**

Frequency and annotation of missense mutations occurring at binding residues (either ligand binding or PPI hotspots, see Supplemental Appendix) or non-binding residues of genes with high number of binding residues in **A.** GENIE v14-public and **B.** MSK-IMPACT NSCLC cohort.

See Supplemental Table S5 for the full list of genes. Stars denote significance (q-value <= 0.1) in Fisher’s exact tests. OncoKB groups include all missense mutations, whereas variant effect predictor groups only include VUSs.

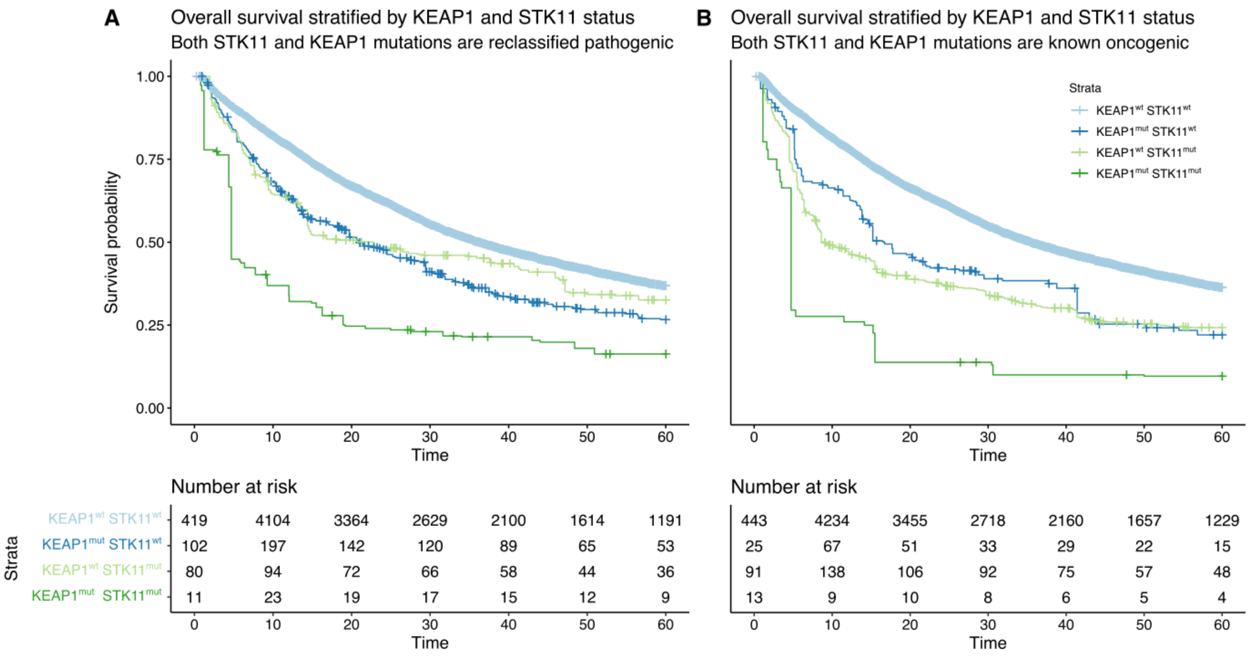


**Figure S9. Outcomes of patients with concurrent *STK11* and *KEAP1* mutations**

A. Kaplan-Meier curves comparing patients with concurrent reclassified pathogenic *KEAP1* and *STK11* mutations versus reclassified pathogenic mutations in either gene or without any mutations.

B. Kaplan-Meier curves comparing patients with concurrent OncoKB oncogenic *KEAP1* and *STK11* mutations versus oncogenic mutations in either gene or without any mutations.

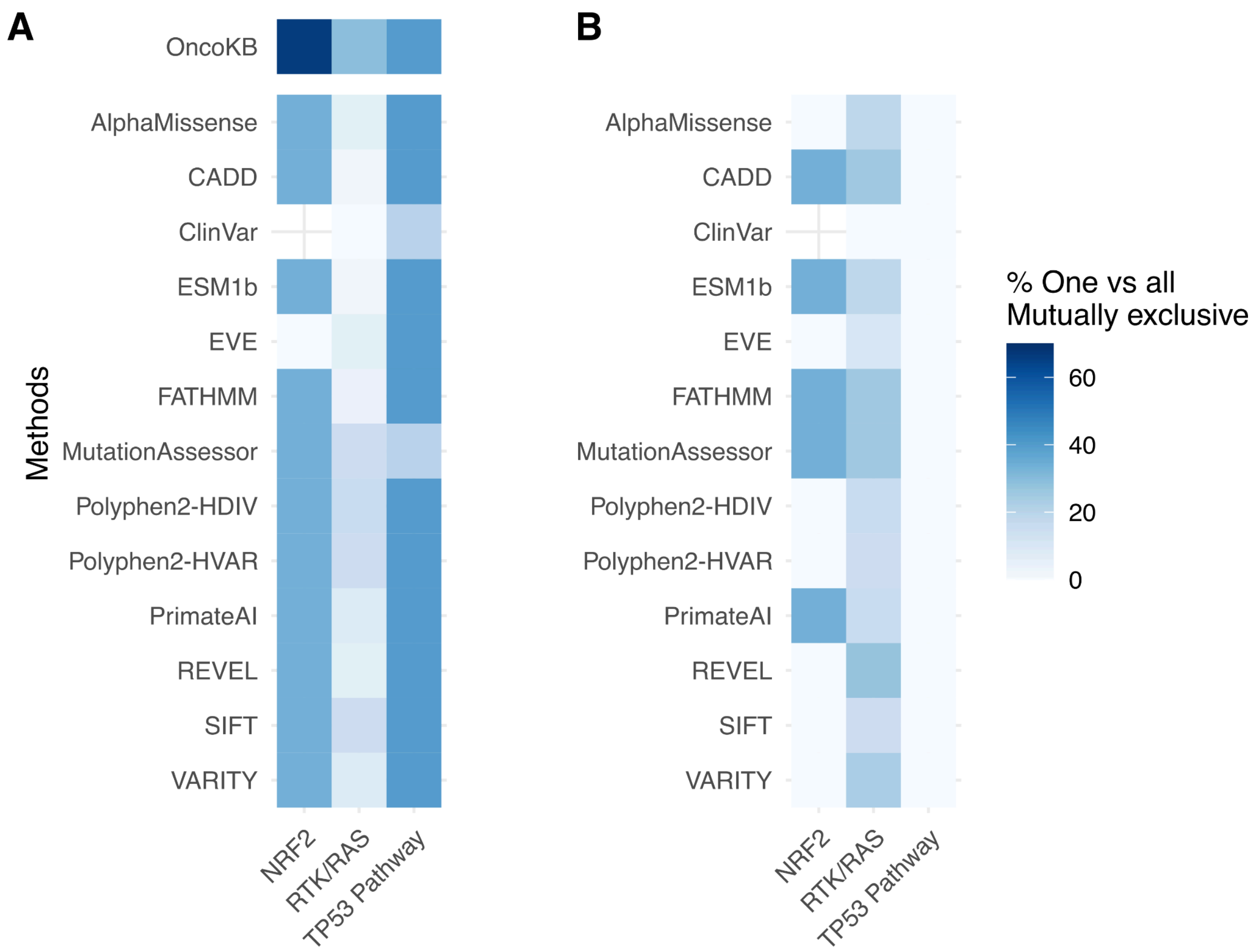


**Figure S10. Mutual exclusivity of mutations in oncogenic signaling pathways**

One-versus-all mutual exclusivity for each gene in a pathway is determined using a two-sided Fisher's exact test. Mutual exclusivity percent for each pathway is then calculated as the number of FDR-corrected significant tests divided by the total number of tests.

A. One-versus-all mutual exclusivity between reclassified pathogenic mutations of each gene and known oncogenic mutations in all other genes in a given pathway.

B. One-versus-all mutual exclusivity between reclassified benign mutations/VUSs of each gene and known oncogenic mutations in all other genes in a given pathway.

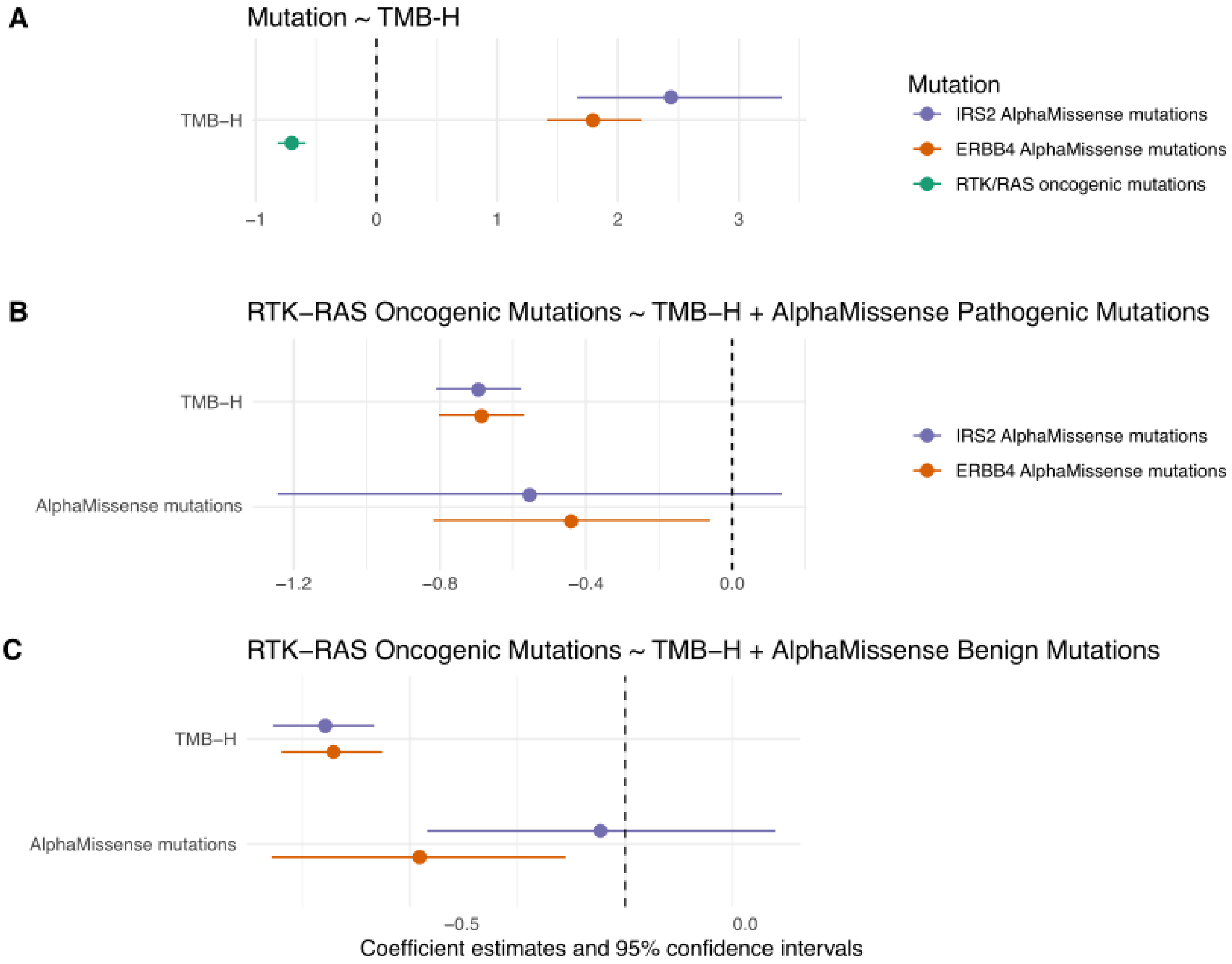


**Figure S11. Mutations in *ERBB4* and *IRS2* are mutually exclusive with RTK/RAS oncogenic mutations independent of TMB-H status**

A. Coefficients for TMB-H status regressed on either RTK/RAS oncogenic mutations or AlphaMissense reclassified mutations in *ERBB4* and *IRS2*. AlphaMissense reclassified pathogenic mutations in *ERBB4* and *IRS2* are positively correlated with TMB-H status, suggesting they occur more frequently in TMB-H samples, whereas RTK/RAS known oncogenic mutations are observed frequently in TMB-low samples.
B. Coefficients for TMB-H status and AlphaMissense pathogenic mutations in either *IRS2* or *ERBB4* regressed on RTK/RAS oncogenic mutations.
C. Coefficients for TMB-H status and AlphaMissense benign mutations in either *IRS2* or *ERBB4* regressed on RTK/RAS oncogenic mutations.

All AlphaMissense mutations in *ERBB4* are significantly and negatively associated with RTK/RAS oncogenic mutations independently from TMB-H status, while all AlphaMissense mutations in *IRS2* are trending towards a negative association with RTK/RAS oncogenic mutations, although small sample sizes in both genes preclude definitive conclusion.

#### Supplementary Tables

**Table S1. Annotation methods overview.** Six variant function prediction methods and two databases were evaluated in this study.

| Method | Description | Required human-curated data |
|---|---|---|
| AlphaMissense | Deep neural network based on the protein structure predictor AlphaFold2, trained on weak labels derived from human and primate germline variants | No |
| CADD | Support vector machine model combining conservation metrics and functional genomic data, trained on weak labels derived from human germline data variants | No |
| ClinVar | Database of human variants and supporting evidences for their functions | Yes |
| ESM1b | Unsupervised deep protein language model | No |
| EVE | Unsupervised deep learning model based on distribution of sequence variation across organisms | No |
| FATHMM | Variant effect predictor based on sequence conservation within hidden Markov models, weighted for cancer-associated mutations | Yes |
| MutationAssessor | Variant functional impact predictor based on evolutionary conservation patterns across protein families and subfamilies | No |
| OncoKB | FDA-approved somatic mutation database | Yes |
| Polyphen2-HDIV | Damaging mutation classifier based on 11 sequence, homology and structural features, trained on Mendelian disease variants and closely related homologs | Yes |
| Polyphen2-HVAR | Damaging mutation classifier based on 11 sequence, homology and structural features, trained on disease-causing mutations and non-disease-related | Yes |

| | | |
|---|---|---|
| | nsSNPs | |
| PrimateAI | Deep learning network, incorporating information from two sub-networks that predict secondary structure and solvent accessibility, trained on common human variants and primate variation | No |
| REVEL | Random forest-based ensemble method trained on other prediction methods and curated variants | Yes |
| SIFT | Sequence homology-based prediction of amino acid substitution | No |
| VARITY (VARITY_R_LOO score) | XGBoost model incorporating scores from unsupervised VEPs as well as information about protein-protein interaction and protein structure. VARITY_R_LOO is the VARITY model trained on rare ClinVar variants (MAF < 0.5%) and cross-validated using a leave-one-variant-out strategy | Yes |

**Table S2. VEP prediction score cutoffs for variant classifications**

| | **GENIE v14** | **MSK-IMPACT NSCLC** | **Non-MSK GENIE BPC NSCLC** |
|---|---|---|---|
| EVE | Class90 | Class90 | Class90 |
| CADD | 24 | 24 | 23.6 |
| ESM1b | -8.96 | -9.11 | -4.3 |
| PrimateAI | 0.67 | 0.67 | 0.6 |
| REVEL | 0.42 | 0.42 | 0.44 |
| VARITY_R_LOO | 0.48 | 0.48 | 0.66 |

**Table S3. Count and proportion of mutations in GENIE v.14-public grouped by their predicted functions and OncoKB annotations.**

Count and proportion of mutations by predicted and annotated functions are provided at

A. mutation level, where each unique mutation is counted once, and
B. population level, where all occurrences of mutations are counted.

**Table S4. Coefficients and confidence intervals from Tukey's range tests for significant difference between predicted scores of non-oncogenic vs. oncogenic mutations.**

Pairwise differences in means of predicted scores between groups were tested for significance using Tukey's range test and p-values were corrected for multiple hypothesis testing using the Benjamini-Hochberg method.

A. Tukey's tests for difference in mean predicted scores at the mutation level.
B. Tukey's tests for difference in mean predicted scores at the population level.

**Table S5. True positive rate at gene level for all missense mutations in GENIE v14-public.**

True positive rate for each gene was calculated as the number of unique missense mutations annotated as oncogenic by OncoKB and pathogenic by a prediction method, divided by the number of total unique missense mutations annotated as oncogenic by OncoKB in each gene.

**Table S6. Reclassified pathogenic mutations occur at ligand-binding residues and protein-protein interaction (PPI) hotspots**

Estimates, confidence intervals and p-values from Fisher's exact tests comparing the enrichment of mutations occurring at binding residues vs. non-binding residues in

A. GENIE v14-public cohort,
B. MSK-IMPACT NSCLC cohort

for being reclassified as pathogenic by different methods.

**Table S7. Cox's proportional hazard coefficients and confidence intervals**

A. Overall survival hazard ratios and 95% confidence interval from weighted Cox proportional hazard models comparing patients with reclassified pathogenic mutations compared to patients without mutation in a given gene (Figure 2D).
B. Overall survival hazard ratios and 95% confidence interval from weighted Cox proportional hazard models comparing patients with reclassified pathogenic mutations compared to patients with oncogenic mutations in a given gene (Figure S6A) and reclassified benign mutations compared to patients without mutations in a given gene (Figure S6B).

P-values were adjusted for multiple hypothesis testing using the Benjamini-Hochberg method.

**Table S8. One-versus-all mutual exclusivity of mutations in oncogenic signaling pathways test results**

Coefficients, p-values and confidence intervals from two-sided Fisher's exact tests used to determine one-versus-all mutual exclusivity for each gene in a pathway. Mutual exclusivity percent for each pathway, shown in Figure S7, is calculated as the number of FDR-corrected significant tests divided by the total number of tests.

A. One-versus-all mutual exclusivity between reclassified pathogenic mutations of each gene and known oncogenic mutations in all genes in a given pathway.
B. One-versus-all mutual exclusivity between reclassified benign mutations/VUSs of each gene and known oncogenic mutations in all genes in a given pathway.

**Table S9. RTK/RAS mutational pattern and TMB-H status of patients with ERBB4 or IRS2 VUSs**

For each patient in the MSK-IMPACT NSCLC cohort with a VUS in either ERBB4 or IRS2, information about the observed mutation, its annotation by AlphaMissense, as well as a list of RTK/RAS pathway genes with known oncogenic mutations and TMB-H status is provided. Each row in the table is one patient; mutational data has been aggregated to patient level for those with multiple samples.

**Table S10. Number of patients in GENIE v14-public with each gene sequenced**
